# A machine learning model predicts protein stability of annotated and alternate protein isoforms

**DOI:** 10.64898/2026.07.16.739020

**Authors:** Océane Marescal, Iain M. Cheeseman

**Affiliations:** Whitehead Institute for Biomedical Research, Cambridge, MA 02142; Department of Biology, Massachusetts Institute of Technology, Cambridge, MA 02142

## Abstract

The regulation of protein stability is essential for cellular homeostasis and is determined by a combination of intrinsic sequence motifs and extrinsic recognition enzymes. Despite growing knowledge of the protein degradation machinery, the ability to predict a protein’s stability from its amino acid sequence remains challenging. Here we develop a machine learning model to predict protein stability from N-terminal amino acid sequences. Using our model and experimental validation, we identify known and novel sequence motifs governing protein stability. We additionally use this model to predict the stability of alternative translational isoforms with distinct N-termini produced from the same mRNA. Despite differing by a limited number of amino acids, we identify N-terminal isoforms with drastically different stabilities relative to their annotated counterparts, highlighting the potential of N-terminal extensions and truncations to regulate protein function. Together, this model provides a valuable tool for evaluating additional protein datasets and protein design strategies.

## Introduction

The total abundance of a protein in the cell is regulated through both its production and turnover, including at the level of protein stability. Protein stability can vary widely between different proteins, with half-lives ranging from minutes for short-lived proteins to years for some long-lived proteins^1,2^. Changes to a protein’s stability can have drastic consequences on both protein and cellular function. Therefore, understanding the rules that govern protein stability is crucial.

A large number of sequence features can affect a protein’s stability, including the identity of the amino acids and their absolute positions within the sequence^3–5^. Sequence or structural motifs within a protein, called degrons, mediate protein degradation through the recognition of substrate proteins by E3 ligases^4,6^. E3 ligases are the final enzyme in the ubiquitination cascade, responsible for the ubiquitination of substrate proteins, a post-translational modification that targets proteins for degradation by the 26S proteasome^7,8^.

Previous work has highlighted the role of protein N-termini in regulating degradation^3,6,9–11^. In a paradigm known as the N-end rule pathway, destabilizing N-terminal residues function as N-terminal degradation signals (N-degrons) and are recognized by E3 ligases for substrate ubiquitination^12,13^. Well-studied N-end rules include the Ac/N-end rule pathway and the Arg/N-end rule pathway. In both of these pathways, the defined sequence determinants for protein degradation are limited to the first two amino acids of the sequence. The Ac/N-end rule pathway recognizes proteins through N-terminal acetylation, which can occur either on the initiator methionine itself or on the second amino acid in cases where the methionine is cleaved cotranslationally by Met-aminopeptidases^3,11,14–16^. For the Arg/N-end rule, protein degradation can be triggered by the presence of a hydrophobic amino acid at the second position following methionine^14^ or by that of certain non-methionine destabilizing residues found at the first position following protein cleavage^3,10^. In contrast to the well-understood influence of the first two residues of a protein on its stability, the effects of amino acids at other positions within the N-terminus on protein stability have not been widely studied.

One manner in which the N-terminal sequence of a protein can differ is through the use of alternative translation initiation sites on a single mRNA^17–21^. Such in-frame alternative start sites can be located upstream or downstream of an annotated start site to produce extended or truncated protein isoforms with distinct N-terminal sequences relative to their annotated counterpart^17,18^. Recent work has highlighted the existence of thousands of N-terminal protein isoforms generated from alternative start sites, with different functions and localizations from their corresponding annotated proteins^18,19^. Given the importance of N-terminal residues on protein stability, such sequence differences at the protein N-terminus could also lead to substantial differences in protein stability.

Here, we develop a machine learning model to predict protein stability from N-terminal amino acid sequences. In combination with experimental validation, we use our model predictions to identify new sequence signatures that affect protein stability. We then use our model to predict the stability of N-terminal extension and truncations and identify protein pairs for which isoform stability differs drastically from that of the corresponding annotated protein.

## Results and Discussion

### Multi-Layer Perceptron predicts N-terminal Protein Stability

To evaluate how N-terminal amino acid composition and sequences influence protein stability, we developed a machine learning model to predict protein stability from N-terminal amino acid sequences (Figure 1A). For model training, we used data from a published large-scale experimental analysis of ∼50,000 peptide sequences and their respective Protein Stability Index (PSI) scores (Timms et al.)^9^. This dataset includes peptide sequences corresponding to the first 23 amino acids of the primary isoforms of all human proteins, with or without the initiator methionine, such that the first residue could be either methionine or another amino acid. PSI scores were determined experimentally using the Global Protein Stability system^9,22,23^ in which an N-terminal amino acid sequence was fused to a GFP to quantify the effect of different peptide sequences on GFP stability based on fluorescence intensity. Scores were continuous values ranging from 1 (most unstable) to 6 (most stable). Using this data, we trained a multi-layer perceptron (MLP) Regressor to predict Protein Stability Index from 23-amino acid peptide sequences (Figure 1A). We used a random split of data for training and testing, with 20 percent reserved for testing, and encoded peptide sequences using one-hot encoding. Our MLP was able to accurately predict PSI scores given N-terminal sequences, with an R^2^ of 0.97 on training data and 0.72 for test data (Figure 1B, C).

**Figure 1:**
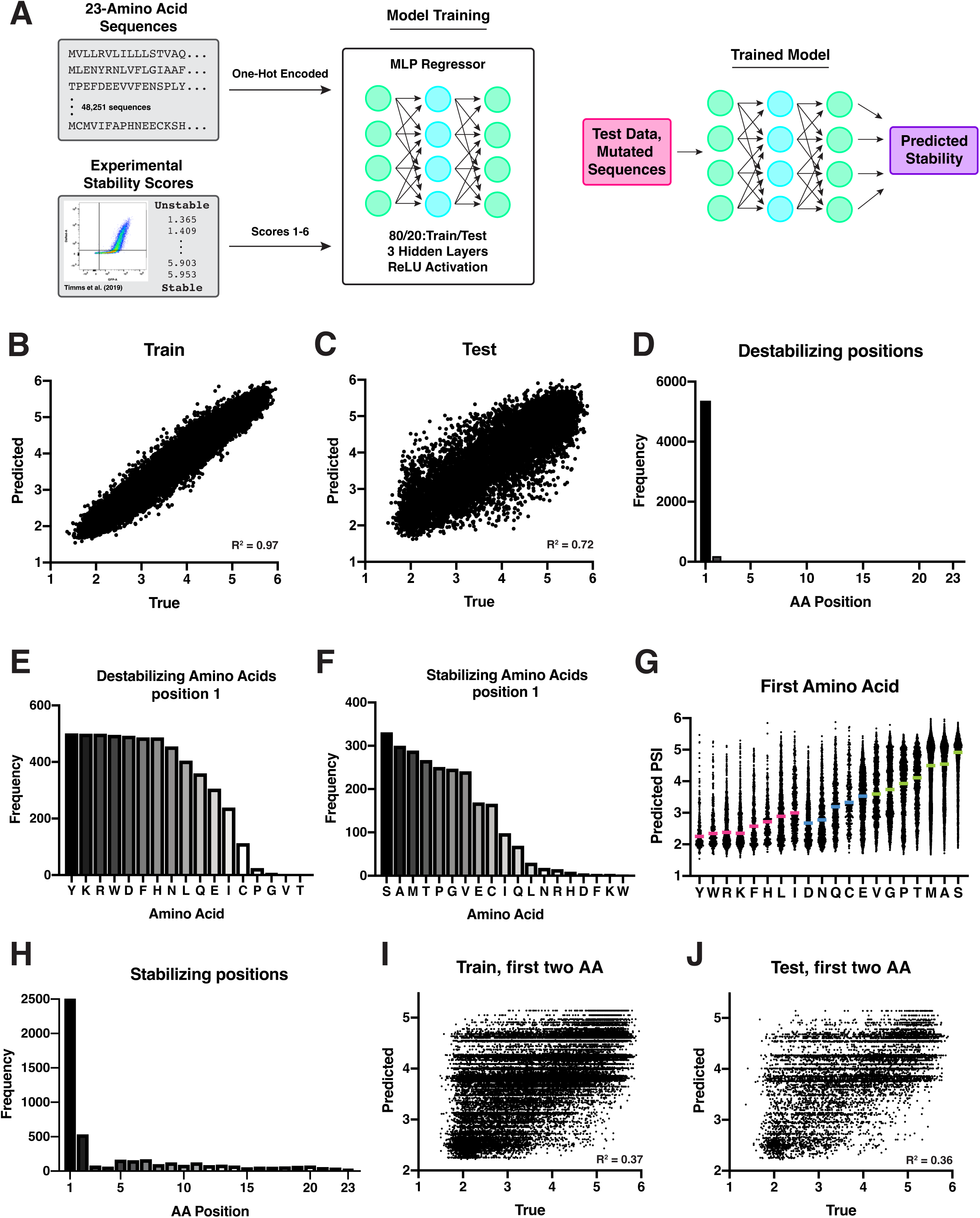
Multi-Layer Perceptron predicts N-terminal Protein Stability. **(A)** Diagram showing MLP Regressor training, input data, and example usage. For more information on model training and parameters, see Materials and Methods section. **(B-C)** Graph plotting the true Protein Stability Index (PSI) (experimentally obtained from Timms et al.) vs. the predicted PSI (predicted by our MLP) for training **(B)** and testing **(C)** data. R^2^ = 0.96 and 0.71, respectively. **(D)** Graph plotting the number of permutated sequences with significantly lower predicted protein stability from the original sequence for each amino acid position changed. **(E-F)** For all sequences where substitution of the first amino acid led to significantly lower **(E)** or higher **(F)** predicted protein stability, this graph plots the identity of what amino acid the first residue was changed to. These indicate amino acids that decrease **(E)** or increase **(F)** protein stability at position 1. **(G)** Graph showing the predicted PSI for all sequences in Timms et al. dataset vs. the identity of the first amino acid in the sequence. **(H)** Graph plotting the number of permutated sequences with significantly higher predicted protein stability from the original sequence for each amino acid position changed. **(J-K)** Graph plotting the true Protein Stability Index (PSI) (experimentally obtained from Timms et al.) vs. the predicted PSI (predicted by our MLP trained on the first 2 amino acids) for training **(J)** and testing **(K)** data. R^2^ = 0.37 and 0.36, respectively.

We next used our trained model to assess the importance of each residue on protein stability. To do so, we perturbed input sequences computationally and assessed the effects of these amino acid changes on the stability scores predicted by our model. We began by identifying residue positions and amino acids that contribute to destabilizing a protein. We selected the top 500 most stable sequences tested by Timms et al. and generated all possible single amino acid permutations of each sequence, substituting each amino acid at each residue for every other amino acid. We then used our MLP to predict the PSI scores of the altered sequences and identified sequences that resulted in significantly lower stability score predictions (decreased stability) relative to the original sequence. We found that replacements in the first amino acid most often led to changes in our model’s PSI predictions (Figure 1D), accounting for 95% of all significantly destabilized permutated sequences. Within the first residue, changes to Tyr, Lys, Arg, Trp, Asp, Phe, His, Asn, Leu, Gln, Glu, and Ile, led to protein destabilization (Figure 1E). These results are in line with prior work on the “N-end rule,” in which destabilizing amino acids at the N-terminus of a protein act as degradation signals^3^. In particular, the destabilizing amino acids identified by our analysis are all established primary Type-I (Arg, Lys, His), primary Type-2 (Leu, Phe, Tyr, Trp, Ile), secondary (Asp, Glu), or tertiary (Asn, Gln) degrons of the Arg/N-end rule pathway^3,24^.

We next conducted a similar analysis to identify residues contributing to protein stabilization. For this analysis, we generated sequence permutations of the top 500 most unstable proteins, and used our MLP predictions to identify stabilized sequences. In this case, changes to the first amino acid contributed to >50% of stabilized sequences. Alterations at the first amino acid that led to increased stability score predictions included Ser, Ala, Met, Thr, Pro, Gly, and Val (Figure 1F, 1H), again confirming prior work demonstrating that the presence of these amino acids with small side chains at the N-terminus promotes protein stability^3^.

Finally, we plotted the PSI predicted by our MLP vs. the first amino acid of the input sequence for all sequences (Figure 1F). Predicted PSI correlated to identities of known N-end rule degrons, with Arg/N-end rule amino acids displaying low PSI (unstable) and known stabilizing amino acids showing high PSI (stable). Thus, our model is able to accurately predict stability based on peptide sequences and can identify known residue positions and amino acid types that previous experimental analyses have determined contribute to protein stability.

### N-terminal protein stability is not fully captured by the first two amino acids

Prior work has focused on the contribution of the first two N-terminal amino acids to protein stability. In contrast, relatively little is known about how other N-terminal protein sequences contribute to overall protein stability. In our permutation stabilization assay, we noticed that changes to other positions downstream of the first two residues also led to differences in predicted stability (Figure 1H). Indeed, 37% of significantly stabilized sequences were caused by single amino acid substitutions present in residues 3-23 (Figure 1H). To determine whether the first two amino acids alone are sufficient to predict protein stability, we trained a second MLP on shortened sequences. We truncated each peptide sequence in the dataset to only its first two amino acids and trained an MLP regressor to predict PSI from these truncated two-mers. If the first two amino acids alone are sufficient to determine a protein’s stability, we would expect a model trained using only the first two amino acids to perform equally well as one trained using the entire 23-amino acid peptide sequence. However, we found that these new models performed far worse than our original models (Figure 1I, J), with an R^2^ of 0.37 and 0.36 on training and test data, respectively. Together, these results indicate that information found in later parts of the sequence also contributes to stability predictions.

### Arginine downstream of the N-terminus may contribute to protein stabilization

To visualize which amino acid substitutions at which residue positions led to increased stabilization of unstable sequences in our permutation assay, we visualized the number of significantly stabilized permutated sequences for each position vs. each amino acid changed in a heat map (Figure 2A, Supplementary Table 1). As expected, substitutions to Met, Ala, Ser, and other known stabilizing amino acids at the first residue resulted in a high number of significantly stabilized sequences (Figure 2A). However, we also found that substitutions of amino acids to Arg or Lys downstream of the N-terminus, from positions ∼4-13, also led to increased predicted stability (Figure 2A).

**Figure 2:**
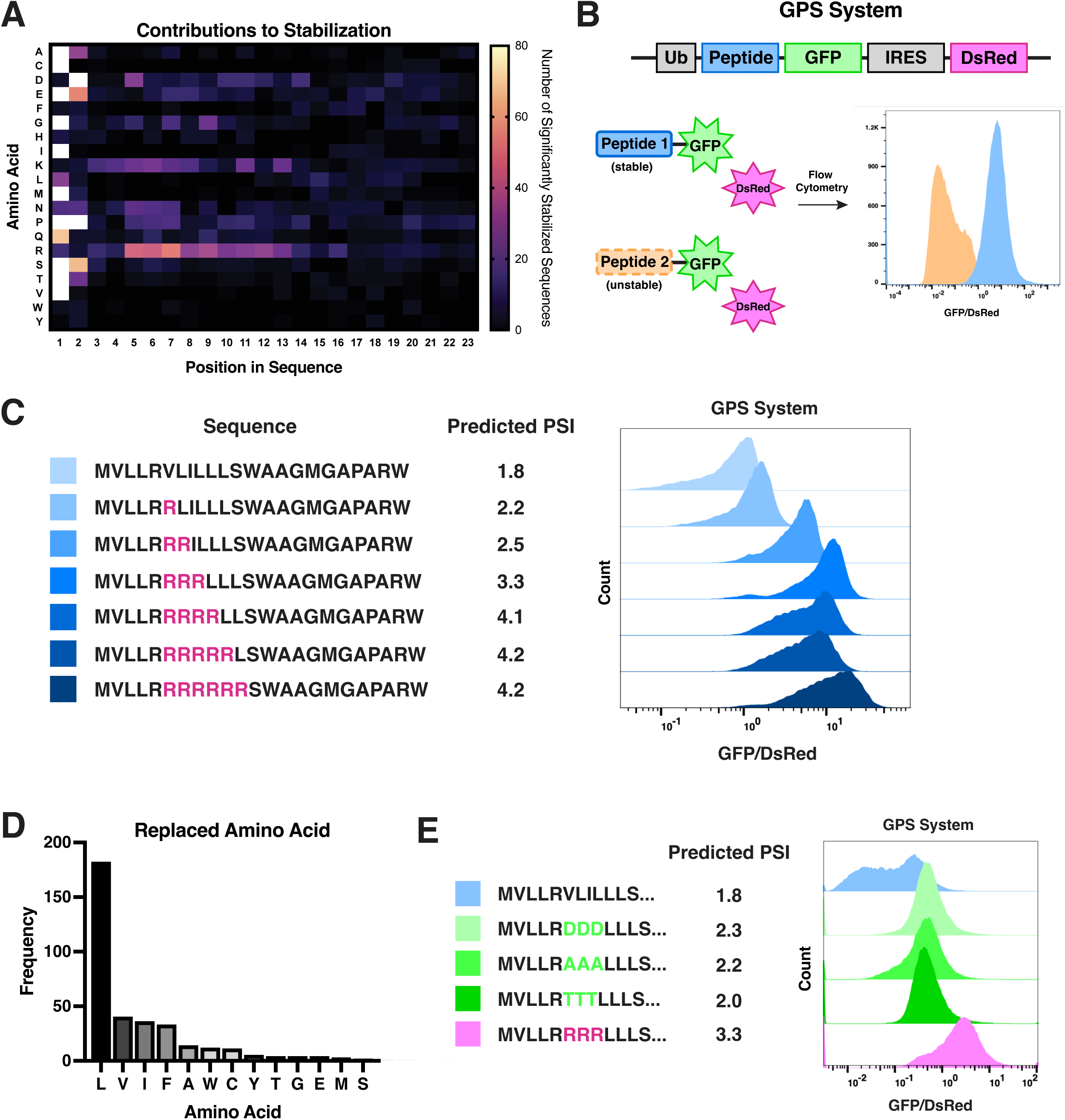
Arginine downstream of the N-terminus may contribute to protein stabilizatio. **(A)** Heat map showing permutation assay results. The number of sequences that were significantly stabilized by position changed (horizontal) and amino acid changed to (vertical). See Materials and methods for more details. **(B)** Diagram showing the Global Protein Stability (GPS) system used to measure the effect of N-terminal sequences on protein stability. An N-terminal peptide sequence of 23 amino acids is fused to a GFP, followed by an internal ribosome entry site and a DsRed as an internal control. Ubiquitin is cotranslationally cleaved by deubiquitinases^3^. After transduction, GFP and DsRed fluorescence intensity are measured in live cells by flow cytometry and the ratio of GFP/DsRed is calculated and plotted. GPS system was adapted from Timms et al. **(C)** Measuring the effect of arginine substitutions at the N-terminus on protein stability. The first 23 amino acids of a known unstable protein, ADAM10, were mutated with increasing numbers of arginines from positions 6 to 11. Left: amino acid sequences. Middle: Protein Stability Index predicted by our MLP Regressor. Right: experimentally determined stability via GPS system (from Figure 2B). Graph shows flow cytometry histogram for each construct, increasing GFP/DsRed ratio indicates increasing protein stability. **(D)** For all sequences that resulted in significantly increased stability upon substitution of an amino acid with arginine, graph plots the identity of the amino acid being replaced by arginine. Amino acids with frequency of only one (D, P, H, N) are not shown for space reasons **(E)** Measuring the effect of arginine substitutions vs. substitutions with other amino acids. The first 23 amino acids of a known unstable protein, ADAM10, were mutated. Positions 6-8 of ADAM10 were mutated to either aspartic acid (DDD), alanine (AAA), threonine (TTT), or arginine (RRR). Left: beginning of the ADAM10 sequence. Middle: Protein Stability Index predicted by our MLP Regressor. Right: experimentally determined stability via GPS system (from Figure 2B). Graph shows flow cytometry histogram for each construct, increasing GFP/DsRed ratio indicates increasing protein stability.

To test whether the presence of Arg at these positions could contribute to protein stability, we designed a series of reporter constructs (Figure 2B, C). Beginning with the two most unstable N-terminal sequences in the Timms et al. dataset (ADAM10, experimental PSI: 1.49 and ZNF208, experimental PSI: 1.41), we generated 6 modified peptides for each with increasing numbers of arginines at positions 6-11 (Figure 2C; Supplementary Figure 1A). The MLP regressor predicted an increase in PSI scores for our designed peptides corresponding to the increasing number of arginine substitutions (Figure 2C; Supplementary Figure 1A). We then experimentally tested the effects of these peptides on protein stability using the established Global Protein Stability system^9,22,23^ (GPS), in which the peptide is placed at the N-terminus of GFP and GFP stability is quantified in cultured human cells by flow cytometry relative to an internal control (Figure 2B). We found that experimentally determined stabilities closely matched our model predictions (Figure 2C, Supplementary Figure 1A). Whereas the original peptide sequences highly destabilized GFP, increasing the number of substitutions of internal N-terminal residues to arginine led to increased GFP stability (Figure 2C, Supplementary Figure 1A).

We next tested whether stabilization occurred due to the replacement of specific amino acids from the original sequence with arginine. For all permutated peptide sequences that were predicted to be significantly stabilized upon the substitution of an amino acid with arginine, we determined the identity of the replaced amino acid (Figure 2D). We found that the vast majority of replaced amino acids were hydrophobic, such as leucine (51%), valine (11%), isoleucine (10%), phenylalanine (9%), and alanine (4%). Indeed, these Arg-stabilized sequences are noticeably enriched for hydrophobic residues from positions 3-13 of the peptides (Supplementary Figure 1B, Supplementary Table 1). An enrichment of hydrophobic residues amongst unstable peptides was also noted in Timms et al.^9^

Increased stabilization resulting from arginine substitutions in these sequences could be due to the loss of destabilizing hydrophobic residues or from the stabilizing presence of arginine itself. Arginine (and lysine) permutations specifically led to the largest number of stabilized sequences in our previous assay (Figure 2A), suggesting that these residues make a distinct contribution to stability beyond simple loss of Leu/Val/Ile. To test this possibility, we computationally compared the effects of Asp and Arg replacements on the predicted stability of hydrophobic-rich N-terminal sequences (Supplementary Figure 2C). For all sequences predicted to be significantly stabilized by arginine replacement in residues 4-14 (Figure 2A), we instead made an aspartic acid substitution (Supplementary Figure 1C). We then used our MLP regressor to predict the stability of the original sequence, the Arg-permutated sequence, and the Asp-permutated sequence (Supplementary Figure 1C). Although replacement of hydrophobic residues with aspartic acid also resulted in increased predicted stability, arginine replacement had a larger predicted stabilizing effect on almost all peptide sequences tested (86%) (Supplementary Figure 1C).

These analyses suggest that the replacement of hydrophobic residues specifically with arginine may have a greater stabilizing effect than their replacement with other amino acids. To test this experimentally, we created 5 versions of our ADAM10-N-terminus-GPS construct: the wild type sequence and 4 mutated sequences (Figure 2C, E). The wild type ADAM10 N-terminus contains a high proportion of hydrophobic residues and highly destabilizes GFP (Figure 2C, E). For our mutated constructs, we compared arginine substitutions within the ADAM10 N-terminus with aspartic acid, alanine, or threonine substitutions. Specifically, we mutated positions 5-7 of ADAM10 (VLI in wild type) with either aspartic acid (DDD), alanine (AAA), threonine (TTT), or arginine (RRR). We then tested the effects of these mutations on protein stability using the GPS system. Replacing VLI with DDD, AAA, and TTT all led to increased predicted and experimental protein stability (Figure 2E). However, RRR-substituted constructs showed an even higher GFP stability (Figure 2E), suggesting an additional stabilizing effect for arginine. Thus, the influence of N-terminal arginines on protein stabilization arises from both the loss of hydrophobic residues and from the presence of arginine itself.

### N-terminal translational isoforms may radically change protein stability

Our model and experimental evaluation confirm that changes to the N-terminal amino acid sequence of a protein can have a strong impact on its relative stability. Recent work using translation initiation site profiling has identified thousands of alternative protein isoforms with distinct N-termini that result from the translation of a single mRNA into multiple, in-frame protein products^18,19,21^. Such alternative isoforms include translational extensions arising from in-frame start sites upstream of the annotated protein sequence and translational truncations generated from downstream start sites (Figure 3A). Given the importance of the N-terminus to overall protein stability, even short extensions or truncations that modify the N-terminal amino acid sequence have the potential to change a protein’s stability.

**Figure 3:**
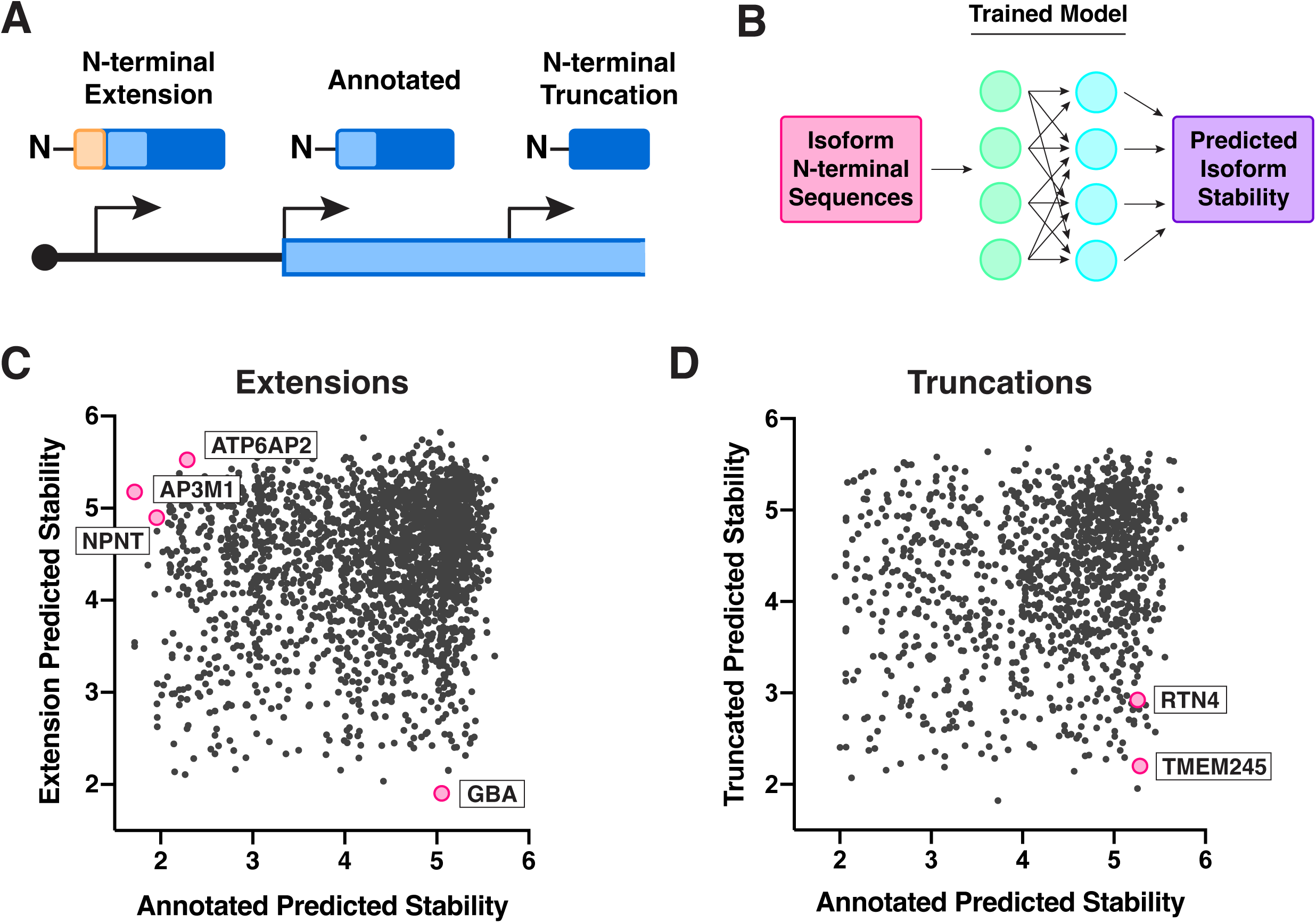
N-terminal translational isoforms may radically change protein stability. **(A)** Diagram showing alternative translation start site isoforms. Start sites can be used upstream or downstream of the annotated start site, resulting in either N-terminal extension or truncation protein isoforms. **(B)** Diagram showing model usage to predict isoform stability. (C-D) Graph plotting the predicted PSI for annotated proteins vs. predicted PSI for the corresponding extended **(C)** or truncated **(D)** isoform(s) of the proteins. Proteins that were further experimentally validated are highlighted in pink.

To determine the effect of novel N-terminal extensions and truncations on protein stability, we used our MLP regressor to systematically predict the PSI score of annotated proteins and their corresponding extended or truncated protein isoforms (Figure 3B). We found that most N-termini, both from annotated and alternative isoform sequences, were predicted to be relatively stable, with predicted PSI scores of 4 or greater (Figure 3C, D, Supplementary Table 2). Importantly, we identified a subset of alternate protein isoforms in which the stability of an extension and truncation was predicted to be substantially different from that of the annotated protein (Figure 3C, D).

To validate our model’s predictions, we generated both extension-annotated and truncation-annotated N-terminal peptide pairs and used the GPS system to experimentally evaluate protein stability (Figure 2B, 3C, D, 4A, B). As predicted, the N-termini of translational isoforms had significant effects on protein stability and could either stabilize (Figure 4C, D, E) or destabilize (Figure 4F, G, H) GFP. Although some of these isoforms only differed from their annotated counterparts by a small number of amino acids (Supplementary Figure 2), the changes in their N-terminal sequences led to large differences in both the predicted and actual stability between the protein pairs. In addition, we noted that alternative N-terminal isoforms with vastly different stabilities relative to their annotated equivalents often differed through the addition or masking of N-terminal hydrophobic amino acid-rich sequences (Supplementary Table 2). As noted above, such hydrophobic regions in N-terminal regions are predicted by our model to result in protein destabilization. For example, both NPNT and ATP6AP2 are unstable proteins with highly hydrophobic annotated N-termini, whereas their N-terminal extension isoforms are stable (Supplementary Figure 2). For these proteins, the addition of an extension masks the N-terminal hydrophobic region of the annotated protein, thus increasing its stability. Conversely, the annotated GBA is a stable protein, but its isoform is unstable due to the addition of a hydrophobic N-terminal extension (Supplementary Figure 2). Thus, N-terminal extension and truncation isoforms can differ widely in stability relative to their annotated counterparts, with the potential to play a physiological role through modulating protein stability.

**Figure 4:**
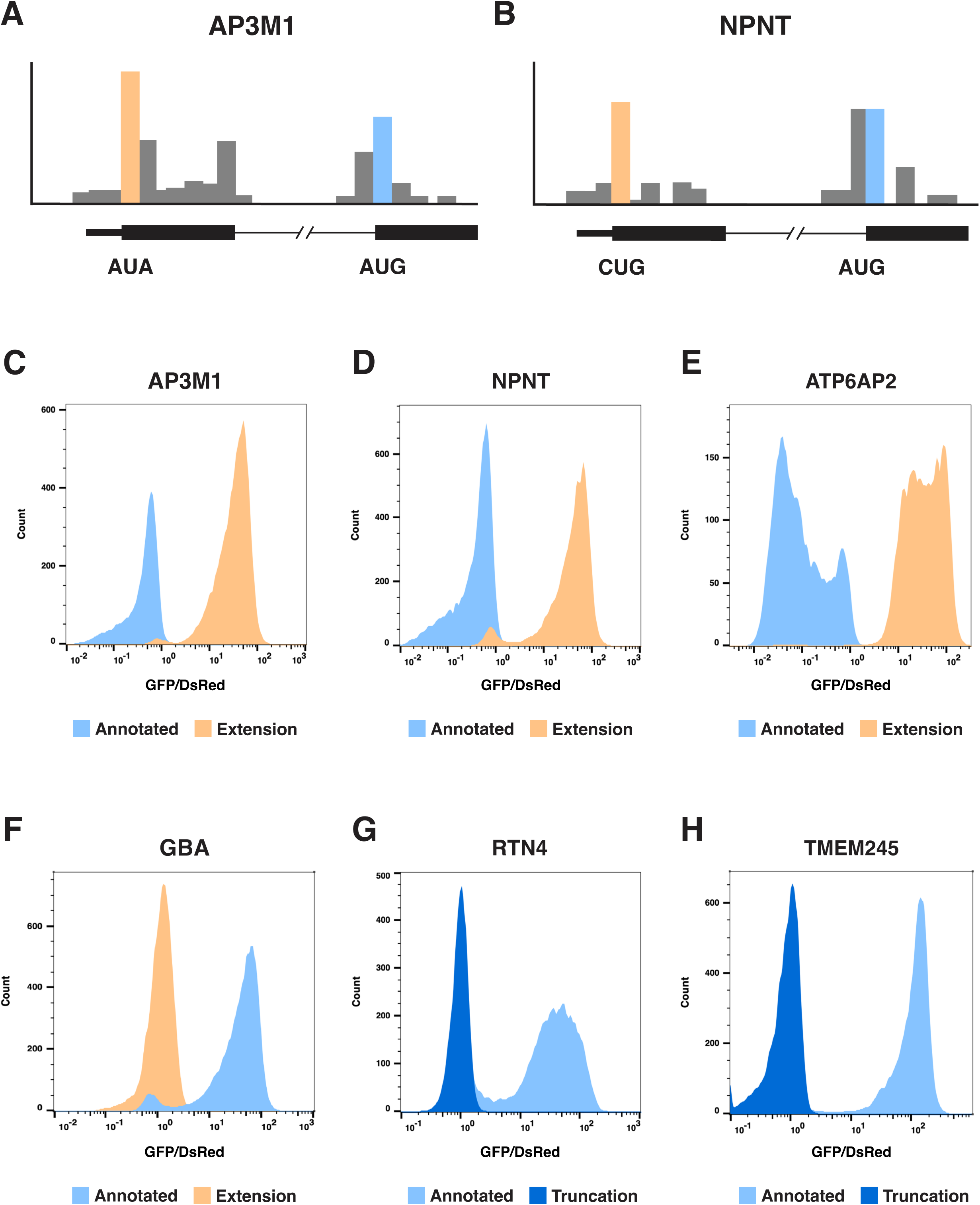
Experimental validation of differential stability of different protein isoforms. **(A-B)** Ribosome profiling traces showing ribosome translation initiation sites and start codons for AP3M1 **(A)** and NPNT **(B)**. See materials and methods for more details. **(C-H)** Experimental determination of the N-terminal stability of different annotated and extension/truncation isoforms from Figure 3 using the Global Protein Stability system. Flow cytometry histograms are shown, increasing GFP/DsRed ratio indicates increasing protein stability. See Materials and Methods for more details.

### A machine learning model for evaluating N-terminal variants

Here, we developed a machine learning model that uses N-terminal amino acid sequences to predict protein stability. We find that the substitution of hydrophobic amino acids for arginine in certain N-terminal peptide sequences increases protein stability. This stability behavior could result from changes in degron recognition and binding by E3 ligases. N-terminal hydrophobic amino acids could act as degrons, which become disrupted upon substitution of these amino acids with arginine. The positive charge of arginine residues could further perturb protein-protein interactions between the E3 and its substrates to a higher extent than substitutions with other amino acids.

We also use our machine learning model to predict the stability of translational isoforms, with many extension and truncation isoforms predicted to have considerably different stabilities relative to their annotated counterparts (Figure 3, 4). This suggests the possibility that cells may use alternative translation to regulate protein abundance. Our previous work found that cells regulate start codon usage in different cellular contexts^18,25^, resulting in altered ratios of alternative isoforms to annotated protein production. Differences in the stability of these isoforms could therefore result in variations in the overall abundance of the proteins depending on cellular context or conditions. Alternatively, cells can also modify translation start-site usage to drive disease through the preferential translation of non-canonical isoforms^26–28^. In this case, the stabilization or destabilization of disease-relevant proteins through alteration of their N-termini could have disastrous consequences for cellular growth or survival. Thus, the modification of a protein’s stability through the use of alternative start sites to produce extension and truncation isoforms with modified N-termini could be a mechanism for the regulation of protein abundance.

The ability to use a machine learning model to predict the stability of N-terminal sequences is a powerful tool for the evaluation of other protein datasets. In addition to translational extensions and truncations, which we have analyzed here, a myriad of regulatory and pathogenic processes can alter the amino acid sequence of a protein at its N-terminus, such as point mutations, alternative splicing, or protease cleavage. Our model can therefore be used to evaluate the stability of all proteins with alternative N-termini, allowing for a deeper understanding of these proteins’ functions.

## Acknowledgements

We thank the members of the Cheeseman lab for feedback throughout the process and the Elledge lab for their generous gift of the GPS plasmid. We also thank Joey Davis and other members of the 6.C51/7.C51 teaching team for their insight and contribution to the machine learning model. This work was supported by grants to I.M.C. from NIGMS (R35GM126930) and the Chan Zuckerberg Initiative (Rare As One). This research was also supported by the Whitehead Innovation Initiative (to I.M.C.).

## Supplementary Figure Legends

**Supplementary Figure 1:**
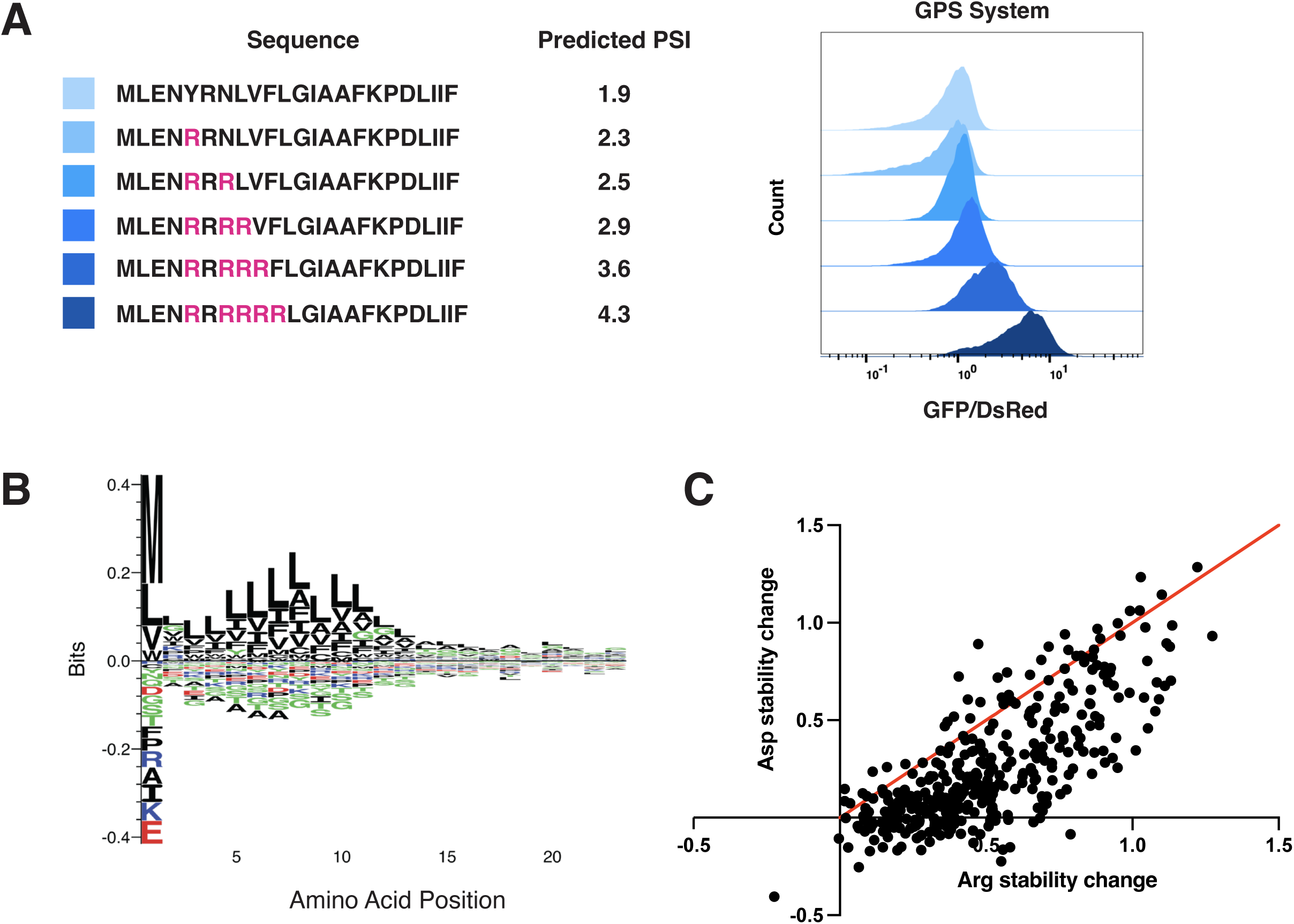
Arginine replacement of hydrophobic amino acids at the N-terminus increases protein stability. **(A)** Measuring the effect of arginine substitutions at the N-terminus on protein stability. The first 23 amino acids of a known unstable protein, ZNF208, were mutated with increasing numbers of arginines from positions 5 to 10. Left: amino acid sequences. Middle: Protein Stability Index predicted by our MLP Regressor. Right: experimentally determined stability via GPS system. Graph shows flow cytometry histogram for each construct, increasing GFP/DsRed ratio indicates increasing protein stability. **(B)** Sequence logo plots for visualization of amino acid enrichment. Plot shows enrichment of hydrophobic sequences in sequences that were stabilized upon arginine substitutions. **(C)** For all sequences that changed stability upon Arg substitution, plots the difference in predicted stability between Arg-substituted (X-axis) and unmutated vs Asp substituted (Y-axis) and unmutated sequences.

**Supplementary Figure 2:**
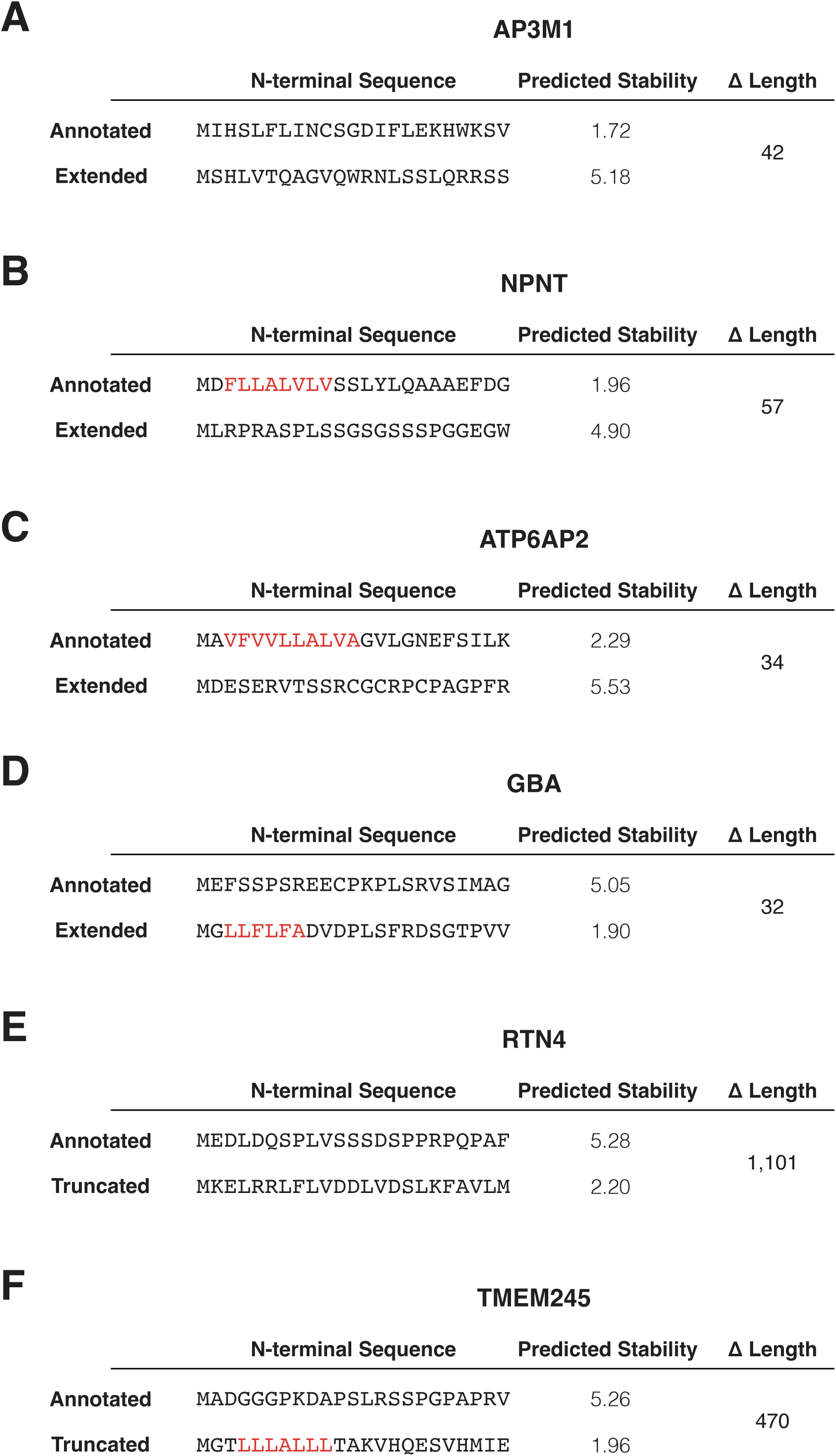
Example of N-terminal isoforms with different stabilities. **(A-F)** Tables showing the N-terminal sequence (first 23 amino acids), predicted stability from our MLP Regressor, and difference in amino acid length for annotated and extended or truncated isoforms of indicated proteins. Hydrophobic sequences are highlighted in red.

**Supplementary Table 1: Stabilizing mutations Tab 1:** Table showing all permutated sequences with significantly increased stability relative to the original sequence, the residue that was substituted, what amino acid it was changed to, and the stability predictions. **Tab 2:** Frequency counts used to make Figure 2A. Shows number of significantly stabilized permutated sequences for each residue position and amino acid changed **Tab 3**: frequency of the amino acids that were replaced by arginine.

**Supplementary Table 2: Extension and Truncation stability predictions** Stability predictions for extension (**Tab 1**) and truncation (**Tab 2**) isoforms and corresponding annotated proteins from our MLP Regressor. Tables show sequences, stability predictions, and calculated differences in predicted stabilities. Highlighted rows show proteins that we followed up on in Figure 4.

## Materials and Methods

### Multi-Layer Perceptron Regressor Design and Training

#### Raw Data

We obtained data from Timms et al.^9^ of 48,251 peptides sequences and their corresponding Protein Stability Index (PSI) scores. Peptide sequences in this paper correspond to the first 23-24 amino acids of the primary isoforms of all human proteins with or without the initiator methionine. Sequences were duplicates, such that those with the initiator methionine were 24 amino acids long and the corresponding sequences without the methionine were 23 amino acids long. PSI scores were determined experimentally using the GPS system and were continuous scores ranging from 1 (unstable) to 6 (stable).

#### Preparation of Input Data

For the peptide sequences, we truncated all 24-amino acid sequences by one residue from the C-terminus, such that all input sequences were 23 amino acids long. Therefore, each sequence was represented twice: once with the initiator methionine followed by 22 amino acids and a second time with 23 amino acids without the start methionine. We then encoded peptide sequences using one-hot encoding, such that the final representation of one peptide was a feature vector of size 460. We used a single random split of data for training and testing, with 20 percent reserved for testing.

#### Model Parameters and Training

Model was designed using scikit-learn. We designed a Multi-Layer Perceptron Regressor. Hyperparameters were: 3 hidden layers of size 512, 256, and 128, ReLU activation function, adam solver, and 0.08 alpha L2 regularization. All other parameters were default from sklearn’s MLPRegressor. Hyperparameters were optimized using hyperopt with r^2^ values on random splits of test data as the objective function to minimize. Hyperparameter search was performed over a small set of hyperparameters with a max evaluation of 10.

#### Model Evaluation

We evaluated the MLP Regressor using r^2^ values and by plotting predicted PSI vs. true PSI for each peptide sequence.

#### First two MLP

For our second MLP trained on only the first two amino acids, we truncated all peptide sequences to the two N-terminal amino acids. For sequences beginning with the initiator methionine, this includes the Met plus one amino acid. For sequences without the initiator methionine, this is the first two amino acids (equivalent of residues 2 and 3 in the methionine-containing equivalents). Sequences were one-hot encoded. PSI scores are the same as used in our first MLP model. We used a train:test split of 80:20. Model parameters are: 3 hidden layers of 512, 256, 128 nodes, ReLU activation, alpha of 0.16, and adam solver. Model was evaluated using r^2^ values, where input data was also truncated 2-amino acid sequences and PSI scores.

### Evaluating contributions to stabilization

We assessed the importance of each amino acid on the prediction by perturbing input sequences. For the top 500 most destabilized and top 500 most stabilized sequences, we generated all possible single amino acid permutations of each sequence and used our MLP to predict the PSI scores of mutated sequences. For each set of mutated sequences derived from one original sequence, we found the standard deviation of the predicted PSIs. We then identified mutated sequences that had significantly different stabilities from the original sequences as those with predicted PSIs +/- 3 standard deviations away from the PSI of the original sequence. We then found what position was changed and the amino acid that it was changed to.

### Predicting Extension and Truncation Stabilities

A list of extensions and truncations and their annotated counterparts was obtained from Ly et al.^18^. We only selected and worked with genes that had only one annotated isoform. However, there can be multiple extended or truncated isoforms for each annotated isoform and we included all isoforms and plotted scores relative to the same annotated protein. The start codon for N-terminal isoforms is sometimes non-canonical (not encoding for methionine), but still is read as methionine upon translation. We converted the first amino acid of these isoforms to methionine in our data to reflect this. Sequences were run through the MLP to give predicted stabilities and then plotted for each protein.

### Statistical Methods and Measures

For R^2^ calculations for our model evaluation (True vs. Predicted for training and test data for our two MLP Regressors), we used sklearn’s MLPRegressor built-in *score* function, which takes one array for predicted values and one array for true values and returns the coefficient of determination. For determining significantly stabilized or destabilized sequences in our permutation assay, we calculated the standard deviation of the predicted scores for permutation sequences of a single original sequence using Python’s NumPy std function and selected sequences that increased or decreased score by 3 calculated standard deviations. (See code for details)

### Code Availability

All code and necessary datasets to run it are available on GitHub: (https://github.com/oceanema/Marescal_and_Cheeseman_2026#).

### Cell culture and Reagents

HeLa cell lines were cultured in Dulbecco’s modified Eagle medium (DMEM) supplemented with 10% fetal bovine serum (FBS), 100 U/ml penicillin and streptomycin, and 2mM L-glutamine at 37°C with 5% CO_2_. Cells were tested routinely for mycoplasma contamination.

For GPS assay, lentivirus was made by transfection with Xtremegene-9 (Roche) of GPS plasmid, VSV-G envelope plasmid, and psPAX2 (Addgene plasmid #12260) packaging plasmid into HEK-293T cells. Cells were transduced with virus in conjunction with polybrene reagent (Millipore Sigma, TR-1003-G).

### Global Protein Stability assay and Flow Cytometry Analyses

The GPS plasmid was a gift from the Elledge lab. N-terminal sequences of interest were cloned using Gibson assembly upstream of the GFP. Cells were transduced with lentivirus containing the GPS plasmid of interest. After 24 hours the media was changed. After another 48 hours, live cells were trypsinized and collected for fluorescence measurements by flow cytometry. For flow cytometry, both GFP and DsRed were measured on a LSRFortessa Flow Cytometer. Flow experiments in Figure 2 and Supplementary Figure 1 were conducted at least 2 times, experiments in Figure 4 were conducted once for each isoform pair. At least 10,000 cells were counted for each condition. Plots were generated using FlowJo V10.10.1 (https://www.flowjo.com/). Histograms show the number of cells (y-axis) vs. GFP/DsRed measurements (Derived Parameter, x-axis) shown on a log scale.

### Recombinant DNA sequences

The following sequences were ordered as gene blocks from Twist Bioscience and cloned into the GPS plasmid (from the Elledge lab) using Gibson assembly between the ubiquitin and the GFP. pOM340-341 were cloned from pOM333 using overlap extension PCR to generate point mutations, the coding sequence only is shown. All plasmids were checked by sequencing. https://docs.google.com/spreadsheets/d/1ZR1ZfhDy7rKyr6R9znI2fAh-9q8c5C108yNy0mAYEFk/edit?usp=sharing

### Ribosome Profiling Traces

Ribosome Profiling data was obtained from Ly et al.^18^ from asynchronous HeLa cells and analyzed using IGV 2.19.8. To generate traces, y-axis (number of reads) was normalized. Ribosome profiling peaks were located 12 base pairs (4 amino acids) upstream of a start site. For clarity, Figure 4A and 4B show the peaks as starting at the start site.

### Sequence logo plots

Sequence logo plots for amino acid enrichment were generated using Seq2Logo-2.0 (https://services.healthtech.dtu.dk/services/Seq2Logo-2.0/). All sequences for which substitution of a residue with arginine led to significant increase in predicted protein stability were used to generate the plot.

## Notes

### Competing Interest Statement

The authors have declared no competing interest.

